# Identification of Direct and Network-Mediated Activation of Retinal Ganglion Cells from Visually Evoked Potentials Using Machine Learning

**DOI:** 10.64898/2026.06.16.732694

**Authors:** Lilli Kiessling, Anna Kochnev Goldstein, Keith Ly, Daniel Palanker

**Affiliations:** Bernstein Center for Computational Neuroscience, Berlin, Germany; Department of Ophthalmology, Stanford University, Stanford, CA, USA; Department of Electrical Engineering, Stanford University, Stanford, CA, USA; Hansen Experimental Physics Laboratory, Stanford University, Stanford, CA, USA

**Author notes:** These authors contributed equally (Lilli Kiessling, Anna Kochnev Goldstein, Keith Ly).

**Keywords:** Prosthetic Vision, Retinal Implant, Neural Selectivity, Visually Evoked Potential (VEP), Synaptic Blockers, PRIMA, Machine Learning (ML), CNN, Integrated Gradients (IG)

## Abstract

**Objective:** To preserve the encoding of visual information in prosthetic vision as close to natural as possible, subretinal photovoltaic implants, which replace the lost photoreceptors, strive to stimulate the second-order retinal neurons, the bipolar cells, while avoiding direct activation of the downstream retinal ganglion cells. To assess the range of such selective subretinal activation, we implanted the devices in rodent models of retinal degeneration and measured the stimulation thresholds based on the visually evoked potentials. After assessment of the bipolar cell-mediated thresholds, direct activation of retinal ganglion cells was measured following intraocular injection of synaptic blockers. Since these chemicals are toxic to the retina, this procedure can only be done once in each animal.

**Approach:** We developed a machine-learning model that identifies the stimulation pathway directly from the recorded visually evoked potentials, eliminating the need for synaptic blockers. The model was trained on recordings from rats implanted with PRIMA subretinal arrays and evaluated on two additional implant architectures, a second rat species, and a different anesthesia protocol.

**Main Results:** The classifier achieved a balanced accuracy of 92% in cross-validation on the training data. Generalization to all unseen experimental conditions yielded an average balanced accuracy of 91%. Integrated Gradients analysis showed that combined bipolar and ganglion cell responses were driven by the early P1 component, while bipolar cell responses relied on later waveform components, consistent with thalamocortical processing dynamics.

**Significance:** The described computational alternative to pharmacological blockers should improve the experimental throughput, allow multiple recordings over the lifetime of the same animal, and might be applicable to optimization of the stimulation settings in patients.

## Introduction

Retinal prostheses aim at restoring vision in patients blinded by retinal degeneration using electrical stimulation of the surviving inner retinal neurons. Subretinal implants replace the lost photoreceptors and target the second-order retinal neurons, the bipolar cells (BCs), as such network-mediated stimulation preserves many features of the natural retinal signal processing [1][2] (Figure 1A). Another approach is a direct activation of the ganglion cells (RGCs) by epiretinal implants. Delivery of the proper retinal code in this case requires identification of many types of RGCs, which represent different aspects of the visual scene [3], and their selective stimulation by corresponding pulse sequencies. Moreover, special care should be taken to avoid stimulation of the RGC axons, which would interfere with the retinotopic mapping of the delivered pattern [4][5][6].

**Figure 1.**
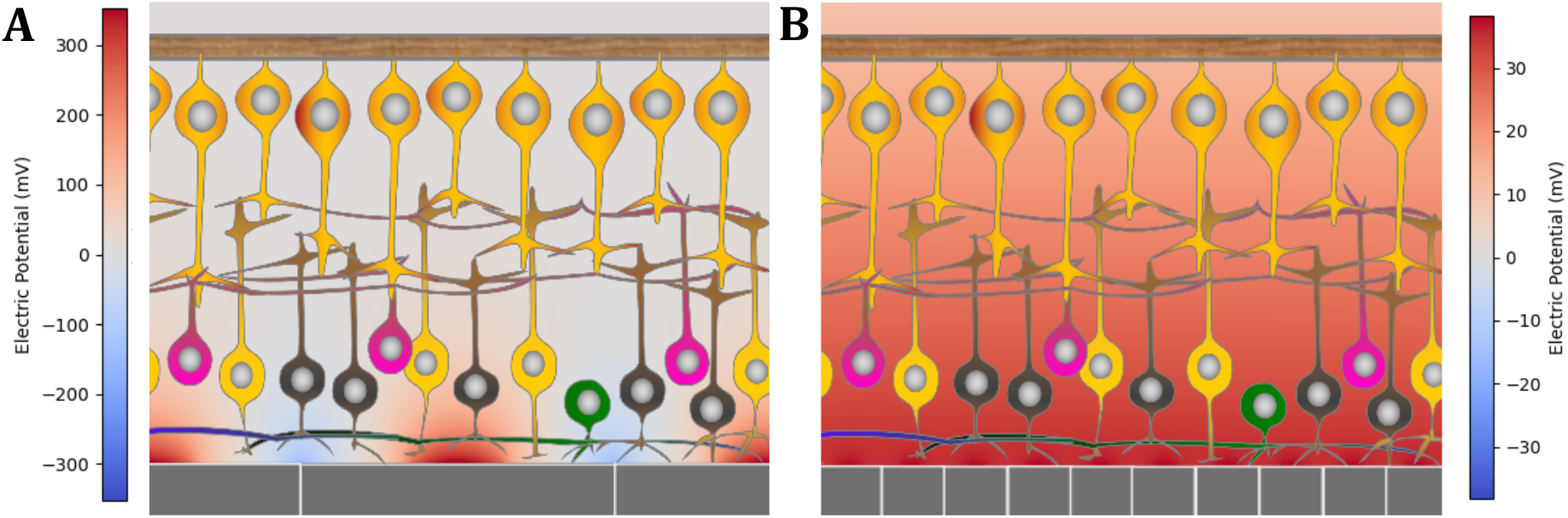
Illustrations of the retinal activation regimes. A. A well-confined electric field that stimulates only the adjacent bipolar cells, thus leading to network-mediated retinal response (PRIMA). B. A poorly confined electric field that activates not only bipolar cells but also directly stimulates ganglion cells (MP20).

Since direct stimulation of RGCs by a subretinal implant would not convey the required neural code and may deteriorate the fidelity of the elicited perception, it should be avoided by proper confinement of the electric field generated by the implant. A poorly confined field penetrates too deep into the retina and increases the chances of direct RGC activation (Figure 1B). Therefore, the distribution of the electric field in the retina should be characterized for each new implant design and each stimulation protocol using computational models and in-vivo testing in animals.

In-vivo characterization of subretinal stimulation by photovoltaic arrays is done by implanting the device in rodents, projecting an infrared light pattern onto the implant, and recording the resulting visually evoked potentials (VEPs) from the animal’s primary visual cortex [7][8][9]. To obtain the strength-duration (SD) curve characterizing the thresholds of a network-mediated retinal response, light pulses are applied at various durations and intensities. For assessment of the direct stimulation of RGCs, synaptic blockers are injected intravitreally into the same animal, which prevents propagation of neural signals from BCs to RGCs. Under such pharmacological isolation, any signals recorded from the visual cortex originate in direct activation of RGCs. The corresponding SD curve serves an upper limit of the network-mediated retinal stimulation with the tested implant [10].

The described procedure discerns the two stimulation pathways, but synaptic blockers are toxic to the retina and other ocular tissues and hence cannot be applied more than once, which limits the experimental throughput, precludes longitudinal follow-up, and increases animal usage. A non-pharmacological method for identifying the stimulation regime based on cortical signals would speed up the device optimization and minimize the use of animals according to the 3R principle (Replacement, Reduction, and Refinement) of ethical animal testing [11].

In rodents, VEP waveforms consist of a sequence of positive and negative peaks (P1, N1, P2, N2, P3, and N3), reflecting successive stages of thalamocortical input and intracortical processing [12][13]. Early components primarily reflect a feedforward geniculocortical drive, whereas later components are increasingly shaped by inhibitory and recurrent cortical dynamics. Direct activation of RGCs introduces unique short-latency components (Figure 2B) and alters the waveform morphology [14][15] compared to BC-mediated retinal activation (Figure 2A).

**Figure 2.**
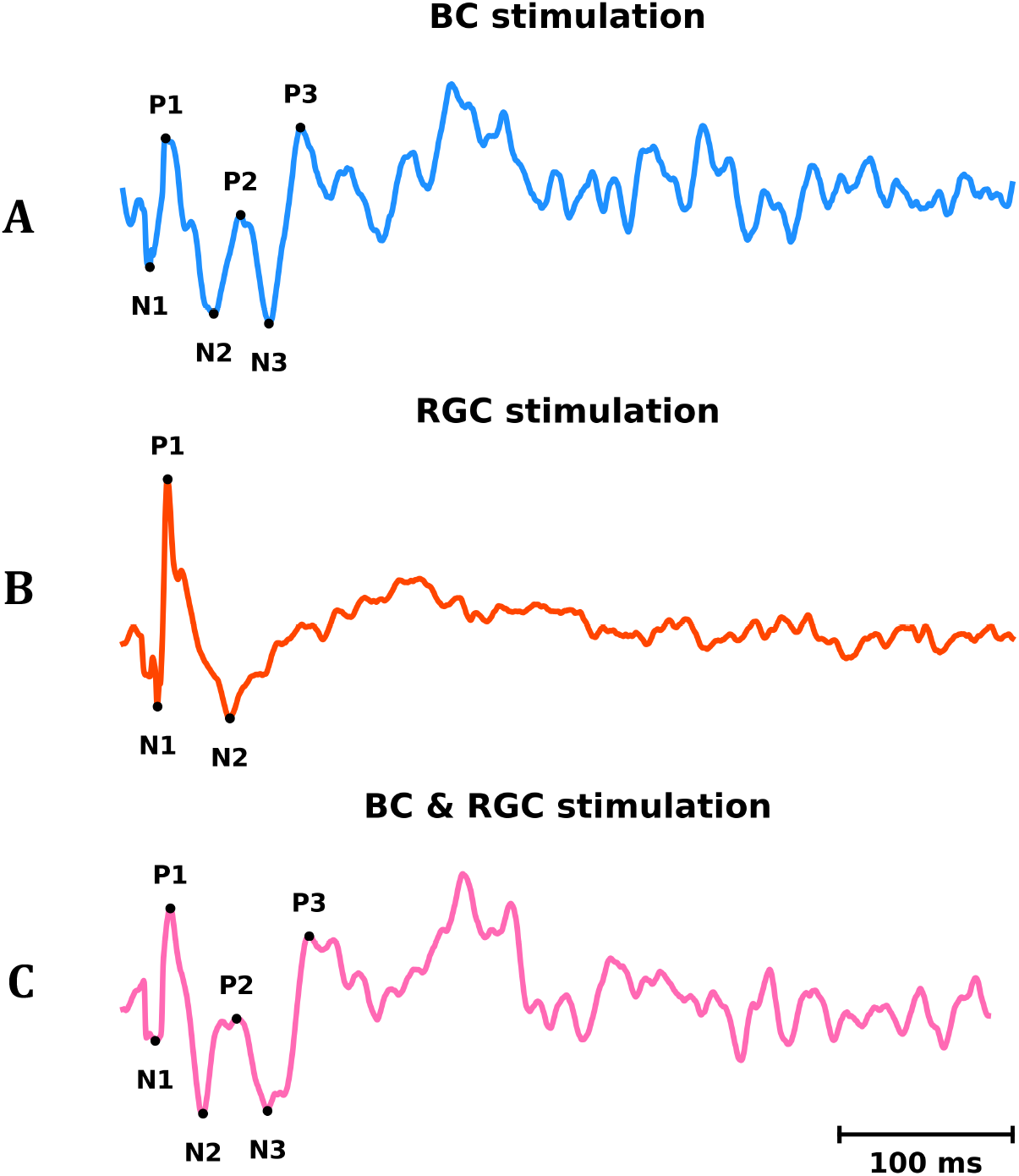
Typical VEP recordings for the three stimulation regimes: **A.** VEP trace measured at low irradiance, when only the network-mediated response is evoked (1.09 mW/mm^2^). **B**. VEP trace measured with synaptic blockers suppressing any network-mediated activity at sufficiently high irradiance to elicit a direct response of RGCs (2.1 mW/mm^2^). **C**. VEP trace measured at high irradiance without synaptic blockers, eliciting both a network-mediated and direct response of RGCs (1.39 mW/mm^2^).

Using the hierarchical temporal structure of VEPs and differences between the two waveform types, we developed a computational framework that infers the stimulation pathway without synaptic blockers - a one-dimensional convolutional neural network (1D-CNN) that was trained to classify VEPs as network-mediated (BC) or combined (BC+RGC) responses.

## Methods

To train the model, VEP recordings from wild type rats implanted subretinally with the PRIMA device were labeled as BC-only and BC+RGC according to the thresholds measured with synaptic blockers. To assess generalizability of the classifier to other experimental conditions, it was then evaluated on various unseen settings, including two other types of implants, a different rat species (with inherited retinal degeneration), and a different anesthesia protocol. To better understand the salient features that drive the classification and provide interpretability, Integrated Gradients (IG) attribution was applied.

### Recording the visually evoked potentials

Following implantation of photovoltaic arrays in rats according to established protocols [7][16], responses to full-field infrared-light (880nm) patterns projected onto the subretinal devices were recorded from the visual cortex. An extensive dataset of VEP traces below and above the threshold for each stimulation regime (BC and BC+RGC) was produced by measurements at multiple laser irradiances and pulse durations, ranging from 0.5 to 10 ms. The light was patterned using a DMD projector illuminated by an 880 nm laser (M1F-880 nm-400 µm, DILAS, Tucson, AZ) and delivered at a repetition rate of 2 Hz. To ensure visibility of the stimulation beam on the implant, a glass coverslip with a viscoelastic material was applied to the cornea, and beam alignment was continuously monitored using a CCD camera.

Rats were anesthetized using two different paradigms – deep and light anesthesia. Both procedures started with an initial dose of 2.5% isoflurane (1 L/min) followed by subcutaneous (light) and intramuscular (deep) administration of ketamine (75 mg/kg) and xylazine (5 mg/kg). For deep anesthesia, this initial dose was supplemented by one quarter of the initial dose given every 15 min during the first hour and every 20 min during the second hour. No maintenance doses were administered for light anesthesia.

VEP signals were recorded using stainless-steel screws placed bilaterally over the primary visual cortex on the rodent skull (4 mm lateral to the midline and 6 mm caudal to bregma). A second screw electrode, positioned 2 mm right of the midline and 2 mm anterior to bregma, served as the reference, and needle electrodes inserted into the nose and tail served as ground. Signals were acquired with the Espion E3 system (Diagnosys LLC, Lowell, MA) at 4 kHz without temporal filtering.

To identify the direct activation threshold of RGCs, when VEP response switched from BC-only to BC+RGC, synaptic transmission was blocked using intravitreal injection of a cocktail of synaptic blockers. The same animal was recorded without and with blockers to obtain the corresponding VEP traces for the two response pathways. Assuming a vitreous volume of 50 µl, the concentrations were 0.8 mM L-AP7, 0.2 mM NBQX, 0.4 mM strychnine, 0.8 mM L-AP4, and 0.32 mM picrotoxin.

### Dataset

To test whether the waveform classification would generalize across various experimental conditions, the VEP dataset was composed of three subretinal prosthetic devices, two rat species, and two anesthesia paradigms. The photovoltaic implants with different extent of the electric field confinement were: (a) PRIMA implant with 100μm pixels in bipolar electrode configuration, where each active electrode is composed of two diodes and surrounded by a local return electrode, yielding a highly confined electric field [17], (b) MP20 implant with 20μm pixels composed of one diode and having a monopolar electrode configuration, where a common return electrode is located along the edge of the implant, resulting in a far-extending electric field [18], and (c) RB20 implant with 20μm pixels composed of one diode and a shunt resistor and having a common return electrode on the back side, producing a field with intermediate confinement in a patterned stimulation [19]. The laser irradiances varied in the range of 0.17–5.97 mW/mm^2^ for PRIMA, 0.08–1.50 mW/mm^2^ for MP20, and 0.12–1.25 mW/mm^2^ for RB20.

Two rodent species were used as an animal model of retinal degeneration: wild type Long–Evans (LE) rats and the Royal College of Surgeons (RCS) rats. In LE rats, local degeneration of photoreceptors above the implant occurs within a few weeks post-op due to blockage of oxygen and nutrient diffusion from the choroid [20]. RCS rats exhibit full degeneration of photoreceptors by six months of age. Lastly, both deep and light anesthesia paradigms were included.

Training of the classifier was done exclusively on PRIMA recordings in LE rats under deep anesthesia (PRIMA LE DA), based on a dataset comprising 203 VEP traces, with 140 BC responses and 63 BC+RGC responses, recorded from 13 animals. All other experimental conditions used for testing the classifier included 128 traces from 4 RCS and 6 LE rats, as listed in Table 1.

**Table 1.**
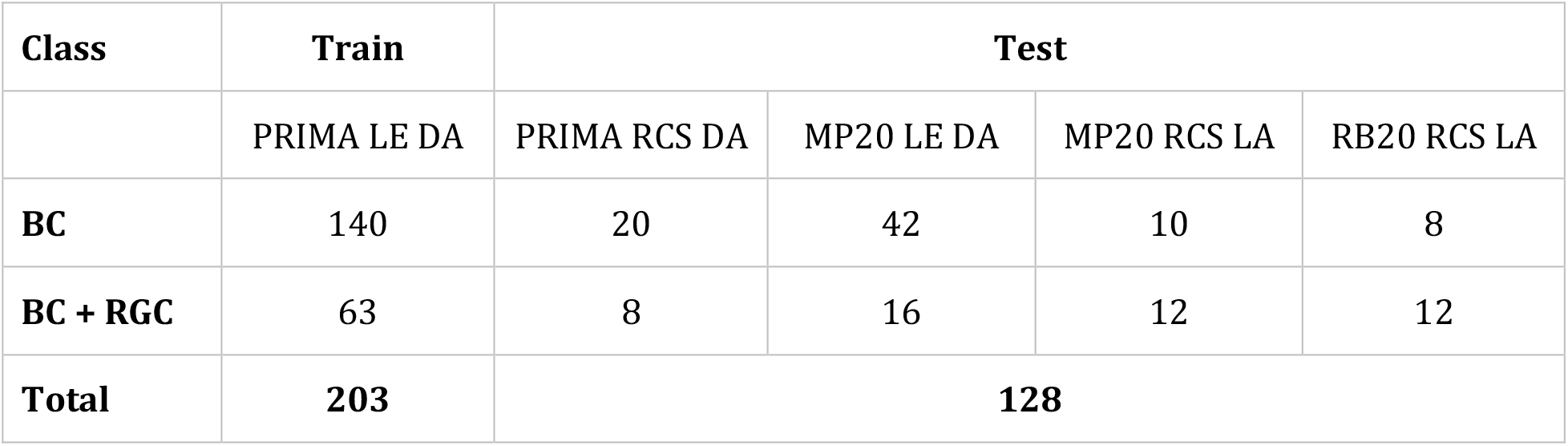
The VEP dataset: data grouped by device (PRIMA, MP20, RB20), rat species (LE vs. RCS), and anesthesia protocol (DA: Deep Anesthesia vs. LA: Light Anesthesia). BC denotes network-mediated response (bipolar cells activation only), while BC+RGC indicates combined BC and direct RGC activation.

The class labels (BC vs. BC+RGC) were assigned based on the RGC activation thresholds measured with synaptic blockers. The response was labeled as BC+RGC if it was measured at an irradiance that exceeded the experimental RGC activation threshold; otherwise, it was labeled as BC. It is important to clarify that blockers were only used to determine the RGC stimulation thresholds, while the VEP traces in the dataset were recorded without blockers.

### Data preprocessing

To ensure standardized inputs, all VEP signals were processed using an automated pipeline. The original VEP recording was an average across 250 trials composed of two consecutive 500ms-long segments acquired under identical conditions. First, the two segments in each recording were averaged to yield a single 500ms-long trace. Next, the quality of the signal was evaluated using a peak-to-peak signal-to-noise ratio (SNR), computed as the ratio between the early post-stimulus response amplitude (0–100ms) and a later baseline window (400–500ms). Six traces with an SNR < 1.0 were excluded.

To ensure equal-length inputs, the remaining signals were truncated to a fixed window. Input dimensionality was reduced while preserving its temporally localized characteristics using discrete wavelet transform (DWT with Daubechies-4, level 4). Only the lowest-frequency approximation coefficients were retained, corresponding to a 16-fold reduction in dimensionality (89 data points per trace). Even though the sampled discrete wavelet transform is not translation-invariant, and hence it induces scale-dependent temporal shifts in the coefficient representation, the relative temporal structure is largely preserved.

Finally, to prevent the classifier from relying on absolute amplitude differences and emphasize the temporal dynamics, each trace was normalized using min-max scaling to the range of [0,1] (Figure 3). Examples of preprocessed VEPs for different irradiances and experimental conditions are shown in Figure 4.

**Figure 3.**
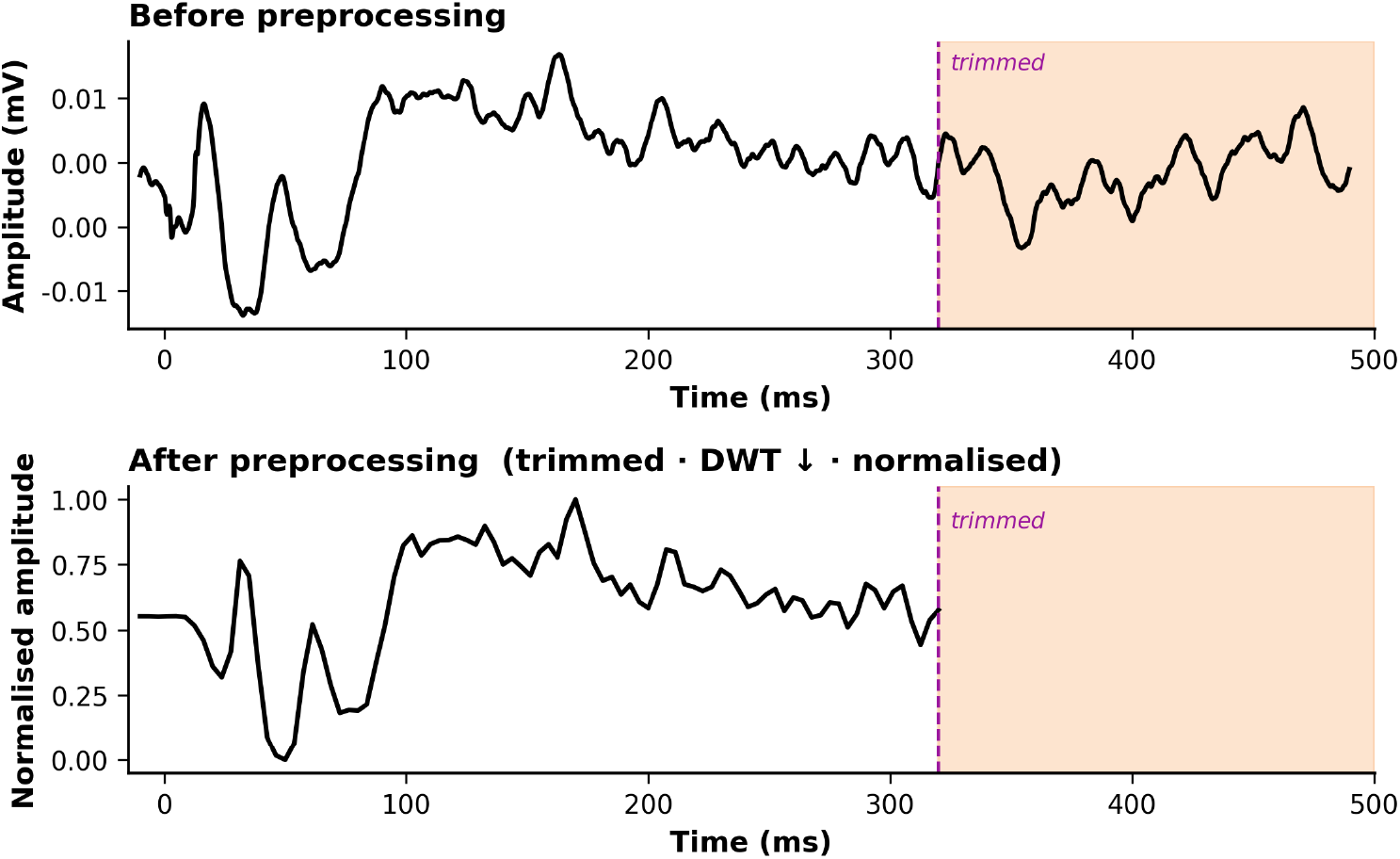
Top: A sample signal averaged over 250 trials and across the two 500ms phases. Bottom: The same signal after checking the SNR, cropping it to 320ms, down-sampling using DWT, and normalizing.

**Figure 4.**
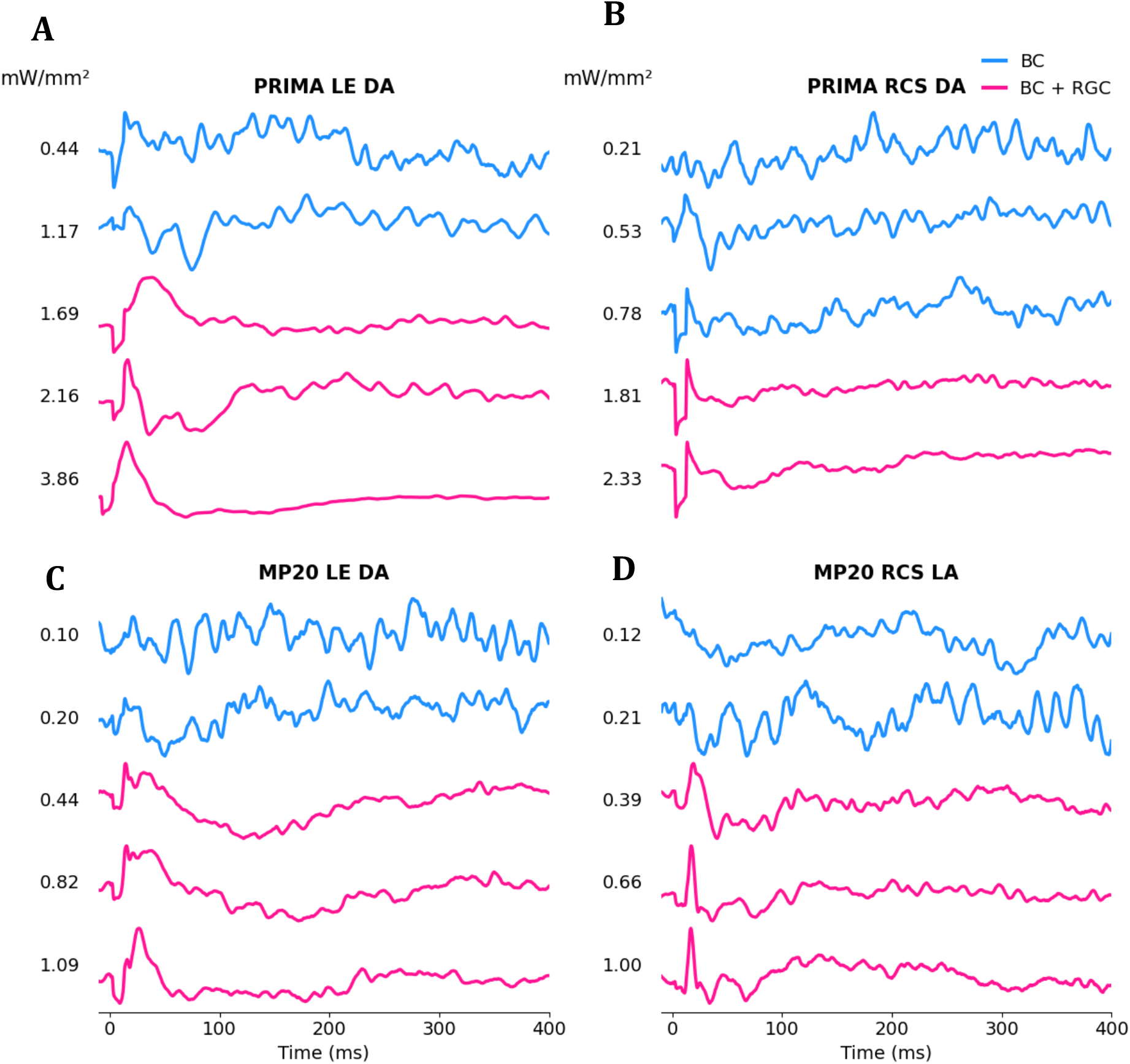
Examples of normalized VEP recordings from a single representative animal at 10ms stimulus duration, across multiple irradiance levels. **A.** PRIMA implant in LE rat under deep anesthesia. **B**. PRIMA implant in RCS rat under deep anesthesia. **C**. MP20 implant in LE rat under deep anesthesia. **D**. MP20 implant in RCS rat under light anesthesia.

### Classification Model

Binary classification of the processed VEP waveforms into BC and BC+RGC regimes was performed using a one-dimensional convolutional neural network (1D-CNN), composed of two convolutional layers followed by a fully connected layer. The first convolutional layer applied 16 temporal filters (with a kernel size of 5 and ReLU activation) with the same padding to preserve temporal resolution, followed by max pooling (with a pool size of 2). The second convolutional layer consisted of 32 temporal filters (with a kernel size of 3 and ReLU activation), followed by subsequent max pooling to further extract the higher-level temporal features while reducing temporal dimensionality. The resulting feature maps were flattened and passed to a dense layer with 32 units (with ReLU activation), followed by a dropout and a softmax layer yielding the class probability.

The network was trained using the Adam optimizer with a learning rate of α = 0.005, batch size of 8, and 30 training epochs. Class imbalance was addressed using a weighted categorical cross-entropy loss, with class weights inversely proportional to class frequencies. To improve generalization, L2 regularization (λ = 10^−4^) was applied to the trainable weights, and dropout (rate 0.2) was used in the fully connected layer. These model hyperparameters were selected based on systematic tuning.

### Integrated Gradients

To interpret the model’s predictions and identify temporally informative signal components, Integrated Gradients (IG) was applied as a post hoc attribution method. IG quantifies the contribution of each input feature by integrating the gradient of the model’s output between a baseline reference signal and the observed input. This approach mitigates the sensitivity of standard gradient-based methods to local nonlinearities and provides more stable and reliable attributions [21].

A zero-valued signal of identical length was used as the baseline, representing the absence of evoked activity after normalization. Attributions were computed for the predicted class and visualized along the temporal axis of the VEP. This analysis enabled the identification of specific latency windows that had a stronger contribution to the classification, thereby providing insights into the physiological features underlying the model’s decisions.

## Results

### Classification Performance

In a 10-fold stratified cross-validation, the 1D-CNN model achieved a balanced accuracy of 92% on the training set consisting of recordings from LE rats with PRIMA implants under deep anesthesia, indicating reliable classification of BC and BC+RGC responses. Class-specific recall reached 95% for BC and 90% for BC+RGC responses, demonstrating balanced sensitivity despite class imbalance. The performance remained high when the classifier was evaluated on independent test sets different from the training set in implant architecture, rat species, and anesthesia protocol. Across the entire external test set, the model achieved a balanced accuracy of 91%, indicating that representations learned during training generalized well to other experimental conditions (Figure 5).

**Figure 5.**
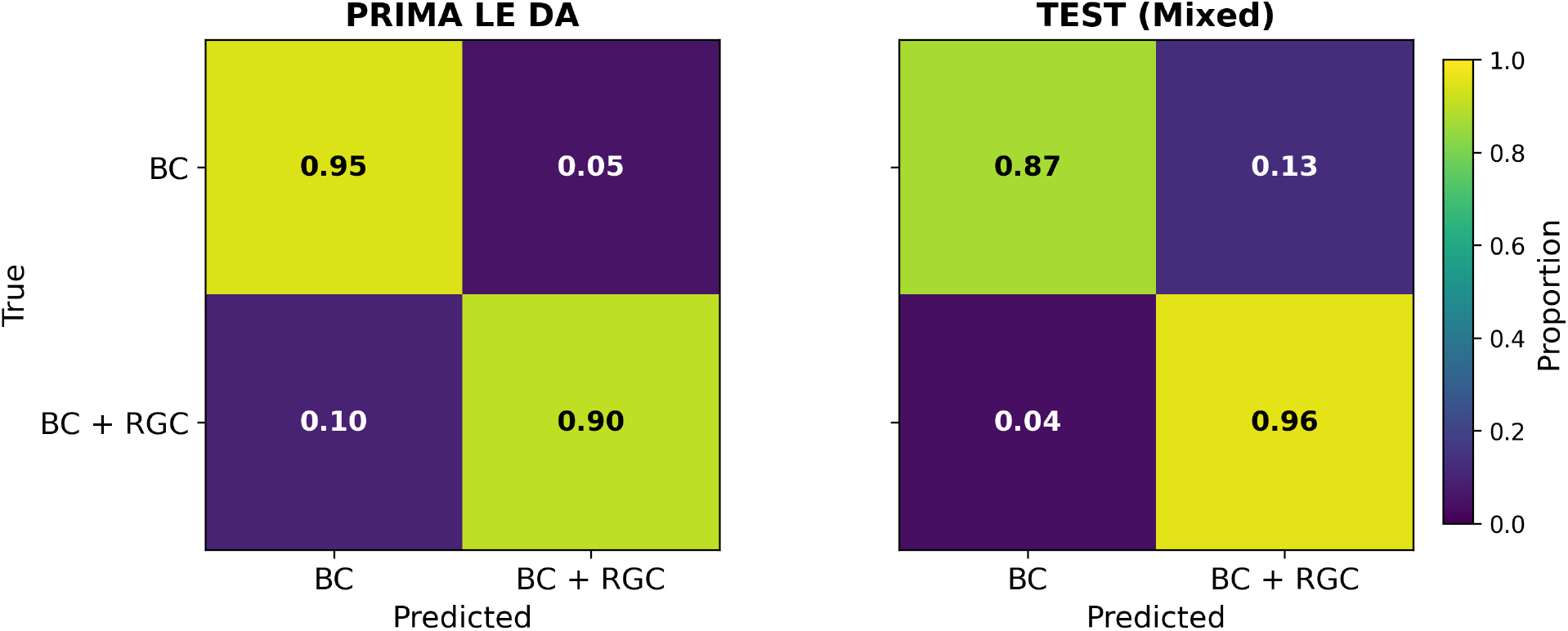
Left: The confusion matrix of a 10-fold cross-validation on LE PRIMA DA recordings. Right: The confusion matrix of evaluating the classifier on all other test conditions.

When analyzed by condition, the performance remained at 90% or above across all conditions. On PRIMA recordings from RCS rats under deep anesthesia, the classifier achieved 90% accuracy across 29 traces, demonstrating that the temporal features learned from LE rats transferred well to a different rat species.

Generalization was also maintained across implant architectures and anesthesia conditions. When evaluated on LE rats with MP20 implants under deep anesthesia, the classifier achieved 90% balanced accuracy across 58 traces. For recordings from RCS rats with MP20 implants and light anesthesia, the classifier achieved 100% balanced accuracy across 20 traces, and for recordings from RCS rats with RB20 implants under light anesthesia, the classifier achieved a balanced accuracy of 96%. In the last two cases the model generalized across a different type of implant, species, and anesthesia, simultaneously. The 100% accuracy obtained in the former case might be due to a smaller sample size for this condition. These results, summarized in Table 2, support the robustness of the learned temporal representation.

**Table 2.** A summary of the balanced accuracy scores achieved for each experimental condition.

| PRIMA LE DA | PRIMA RCS DA | MP20 LE DA | MP20 RCS LA | RB20 RCS LA |
| --- | --- | --- | --- | --- |
| 92% | 90% | 90% | 100% | 96% |

To further assess the performance of the tool for practical deployment, a 10-fold cross-validation was performed on the combined dataset of all measured traces across all experimental scenarios. This validation yielded a balanced accuracy of 92 ± 7% . The performance remained consistent at individual scenarios with balanced accuracies of 91% (PRIMA LE DA), 97% (MP20 LE DA), 94% (PRIMA RCS DA), 96% (MP20 RCS LA), and 85% (RB20 RCS LA). These results demonstrate that each test case can be robustly classified within a unified model despite the differences in implant design (i.e. electric field penetration depth) and other experimental conditions.

### Integrated Gradients Analysis

The Integrated Gradients attribution maps revealed distinct temporal features for each of the two stimulation regimes (Figure 6). For VEP traces classified as BC+RGC, strong attribution was consistently concentrated around the early P1 component. In contrast, VEP traces classified as BC exhibited the strongest attribution at later components, particularly near P3. As individual peaks cannot always be reliably identified by visual inspection alone, manual annotations were restricted to the exemplary PRIMA LE DA traces (Figure 6A). In these examples, the detected components corresponded well to the canonical P1, P2, and P3 structure. In less distinct traces, the classifier relied on broader waveform components, indicating that decisions are based on distributed temporal features.

**Figure 6.**
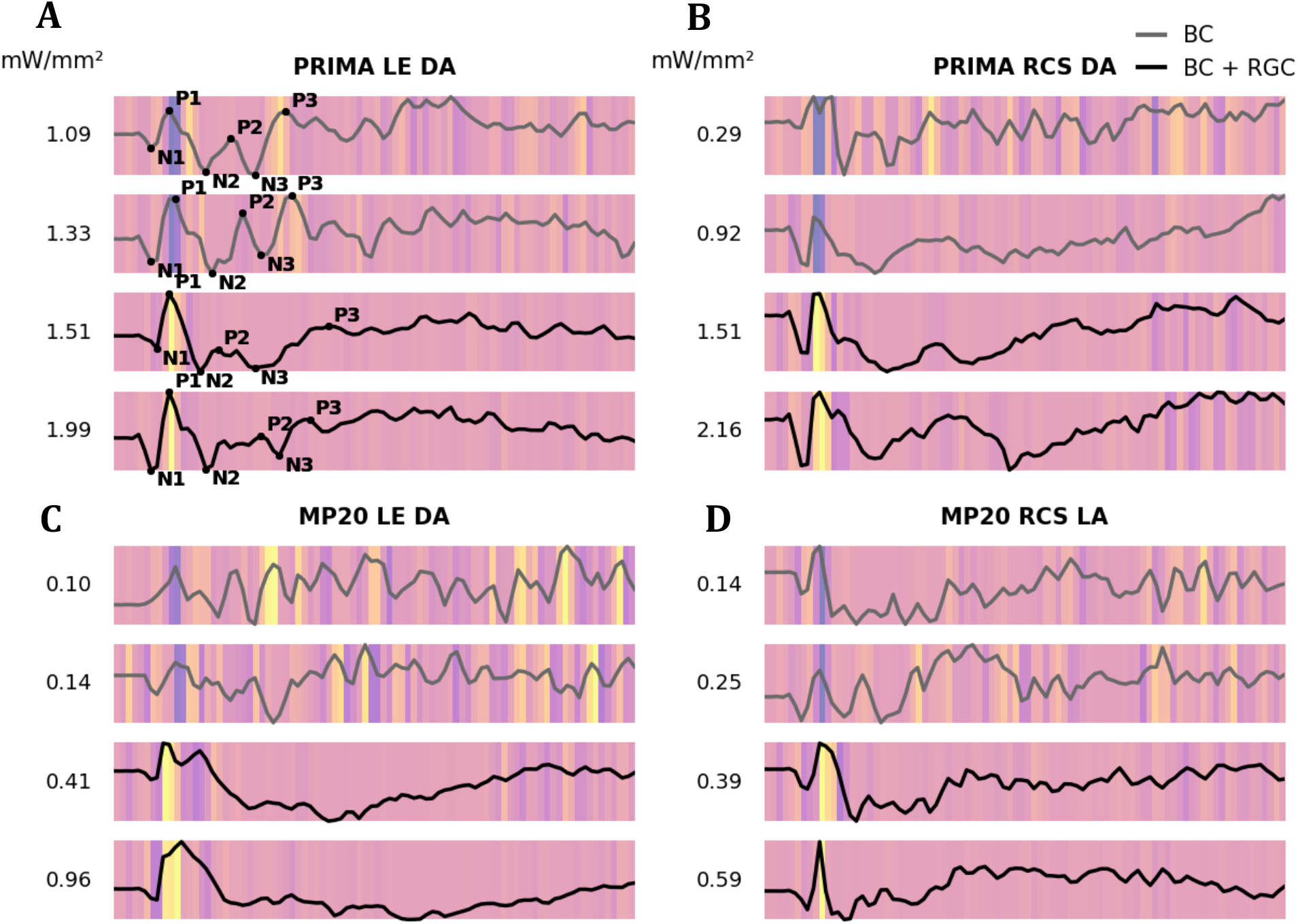
IG attribution maps for representative VEP traces elicited by 10ms pulses across different experimental conditions. Each panel shows sample responses from a single representative animal per condition (different animals across conditions). Yellow indicates features with strong positive attribution, whereas purple indicates negative attribution - signal components the presence or absence of which contributes the most to the classifier’s decision. **A.** PRIMA in LE rat under deep anesthesia. **B**. PRIMA in RCS rat under deep anesthesia, **C**. MP20 in LE rat under deep anesthesia **D**. MP20 in RCS rat under light anesthesia.

To assess whether classification performance depends on signal duration, the model was trained and evaluated on VEP segments of varying lengths, ranging from 20 to 400ms. While performance within the PRIMA LE DA condition remained relatively high (above 80%) even for short input windows, generalization across the independent test conditions improved markedly as longer portions of the waveform were included, with balanced accuracy increasing from approximately 50-60% for 20ms windows to 90–100% for longer windows (Figure 7). Dependence on the signal length shows that classification is not determined solely by the short-latency P1 component but rather by the physiological information distributed across the broader VEP waveform.

**Figure 7.**
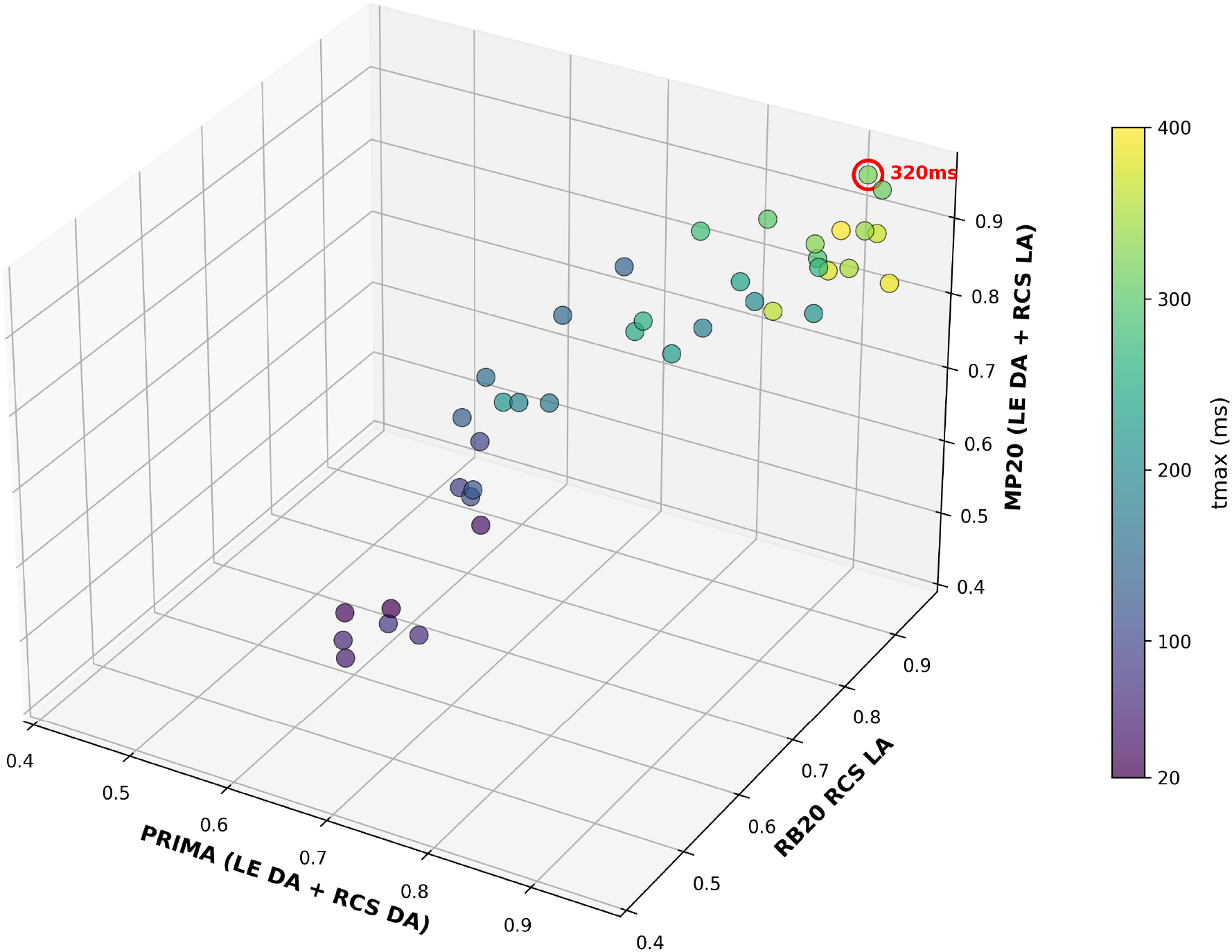
A Pareto plot of the test conditions, grouped by implant type, showing the influence of signal length (the parameter Tmax) on the balanced accuracy scores.

Electrophysiological studies have shown that while the early P1 peak reflects the initial thalamocortical excitation, later components are shaped by inhibitory and recurrent cortical dynamics within the thalamocortical network [22]. The attribution patterns suggest that the classifier relies on physiologically meaningful temporal features. When later components are present, the model reduces the importance of the early peak, indicating that the extended temporal structure in the waveform serves as evidence against direct RGC activation.

## Discussion

The proposed compact 1D-CNN model successfully performed a binary classification of the VEP traces into BC and BC+RGC categories, with a balanced accuracy of above 90%, generalizing across implant architectures, rat species, and anesthesia conditions. This outcome demonstrates that visually evoked potentials contain sufficient information to distinguish between the network-mediated (BC) and combined BC+RGC activation of the retina. The consistently high performance across various experimental conditions emphasizes that this distinction between activation regimes is robustly encoded in the VEP waveforms.

The Integrated Gradients analysis further indicates that the classifier relies on physiologically meaningful components of the VEP waveform. Responses classified as BC+RGC were dominated by the early P1 component, whereas BC responses showed stronger contributions from later components, particularly P3 and beyond. This pattern is consistent with the thalamocortical processing dynamics: early VEP components reflect primarily the feedforward geniculocortical input, while later components arise from recurrent thalamocortical and intracortical processing. Our results indicate that direct RGC activation, which bypasses synaptic processing within the inner retina and therefore arrives earlier to the cortex, is dominated by rapid VEP components. In contrast, network-mediated activation engages retinal circuitry before eliciting ganglion cell spiking, introducing additional temporal structure into the response. Consistently, models trained on shorter input windows showed reduced generalization across experimental conditions, and performance improved when longer temporal segments of the VEP signal were included. These findings suggest that the retinal response is encoded in the full temporal waveform of the VEP signal rather than in a single latency or amplitude metric. The temporal richness of network-mediated prosthetic VEPs parallels the multiphasic waveform structure observed in natural visual responses in rodents, where the full N1-P1-N2-P2 sequence reflects the sequential engagement of photoreceptors, horizontal cells, and bipolar cells, before ganglion cells spiking [21][22].

VEP morphology was also affected by the depth of anesthesia. Ketamine–xylazine anesthesia is known to modify cortical responses in a dose-dependent manner, typically prolonging latencies and altering amplitudes, as observed in rats [25] and in mice [26]. In our studies, VEPs obtained under light anesthesia exhibited richer temporal structure and more pronounced later components, consistent with preservation of more complex cortical dynamics under lighter anesthetic states [27]. The highest performance (96% and 100% classification accuracy) was observed under light anesthesia (RCS rats with MP20 and RB20 implants), indicating that a richer temporal structure likely contributes to stronger separability.

The classifier also generalized well across implant architectures with different electric field distributions. PRIMA has bipolar pixels with local return electrodes whose electric field is well-confined, whereas MP20 and RB20 employ a monopolar configuration with a global return on the edge or on the back of the implant, and its electric field penetrates much deeper into the retina. Despite these differences, the model trained on traces recorded with PRIMA maintained high accuracy on MP20 and RB20 recordings, suggesting that the learned features reflect physiological activation regimes rather than device-specific signal characteristics and therefore should be applicable to various future device architectures, including 3-dimensional electrodes [28].

In conclusion, the presented approach to identify direct RGC activation in prosthetic vision provides a computational alternative to synaptic blocker experiments. This way, the same animal can be tested repeatedly over time, enabling longitudinal assessment of the stimulation thresholds and device performance under various stimulation protocols. This capability should accelerate the evaluation and optimization of retinal prostheses while utilizing the best practices of ethical animal research. It might also be extended to human patients for optimization of the stimulation parameters. Overall, these results provide a proof of concept that machine learning can infer retinal activation pathways directly from cortical signals and offer a practical experimental tool for its implementation.

## Acknowledgements

Studies were supported by the National Institutes of Health (Grants R01-EY-035227, and P30-EY-026877), the Department of Defense (Grant W81XWH-22-1-0933), AFOSR (Grant FA9550-19-1-0402), and an unrestricted grant from the Research to Prevent Blindness.

## Data Availability

This computational framework was developed in Python and is openly available at: https://doi.org/10.5281/zenodo.20708698.

